# Set-Min sketch: a probabilistic map for power-law distributions with application to *k*-mer annotation

**DOI:** 10.1101/2020.11.14.382713

**Authors:** Yoshihiro Shibuya, Djamal Belazzougui, Gregory Kucherov

**Affiliations:** LIGM, CNRS, Univ. Gustave Eiffel, Marne-la-Vallée, France; CAPA, DTISI, Centre de Recherche sur l’Information Scientifique et Technique, Algiers, Algeria; Skolkovo Institute of Science and Technology, Moscow, Russia

**Keywords:** sketching, Set-Min sketch, *k*-mer counting, *k*-mer frequency spectrum, power-law distribution

## Abstract

**Motivation:** In many bioinformatics pipelines, *k*-mer counting is often a required step, with existing methods focusing on optimizing time or memory usage. These methods usually produce very large count tables explicitly representing *k*-mers themselves. Solutions avoiding explicit representation of *k*-mers include Minimal Perfect Hash Functions (MPHFs) or Count-Min sketches. The former is only applicable to static maps not subject to updates, while the latter suffers from potentially very large point-query errors, making it unsuitable when counters are required to be highly accurate.

**Results:** We introduce Set-Min sketch – a sketching technique for representing associative maps inspired by Count-Min sketch – and apply it to the problem of representing *k*-mer count tables. Set-Min is provably more accurate than both Count-Min and Max-Min – an improved variant of Count-Min for static datasets that we define here. We show that Set-Min sketch provides a very low error rate, both in terms of the probability and the size of errors, at the expense of a very moderate memory increase. On the other hand, Set-Min sketches are shown to take up to an order of magnitude less space than MPHF-based solutions, especially for large values of *k*. Space-efficiency of Set-Min takes advantage of the power-law distribution of *k*-mer counts in genomic datasets.

**Availability:** https://github.com/yhhshb/fress

## 1 Introduction

Counting *k*-mer occurrences in genomic sequences is a rather common task in many bioinformatics pipelines. It is often the first step performed before a more complicated analysis, and its main applications go from read trimming [1] to alignment-free variant calling [2, 3]. In recent years, many *k*-mer counting algorithms have been proposed, such as Jellyfish[4], DSK[5] or KMC[6].

All these tools output a map associating *k*-mers to their counts. Such a map can require a fairly big amount of disk space, especially when large values of *k* are used. For example, the KMC output for a human genome with *k* = 32 weights in at around 28 GB. Even with compression, memory efficiency remains an important issue. A way to tackle this problem is to only store counters, without the *k*-mers. This idea is supported by the fact that, in many applications, we only deal with *k*-mers that come from partially assembled reads or succinct representations of *k*-mer sets such as colored de Bruijn graphs[7], or spectrum-preserving string sets [8, 9], that is, only *k*-mers present in the original data are queried for their frequencies.

The idea of storing only counter information and not *k*-mers is also supported by the observation that the number of distinct *k*-mer counts in genomic data is relatively small. It is known that *k*-mer counts in genomes obey a “heavy-tail” power-law distribution^1^ with a relatively large absolute value of the exponent [10, 11]. For such distributions, the number of *distinct k*-mers makes a linear fraction of the data size, while the number of *distinct k*-mer counts is relatively small. For example, for the human genome and *k* = 27, there are about 2.5 billions distinct *k*-mers but only about 8,000 distinct frequency values. Moreover, the majority of *k*-mers have a very small count: in the above example, 97% of *k*-mers are unique and 99% of *k*-mers have a count of at most 5. Frequent *k*-mers often tend to have an identical count as well, due to transposable elements: for example, *k*-mers specific to Alu repeats in primate genomes will likely have the same count.

### Our contribution

We propose a new probabilistic data structure that we call *Set-Min sketch* capable of representing *k*-mer count information in small space and with small errors. The sketch guarantees that the expected *cumulative error* obtained when querying all *k*-mers of the dataset can be bounded by *εN* where *N* is the number of *all k*-mers (i.e. essentially, the size of the dataset). We provide a theoretical analysis in order to dimension the sketch according to the desired error bound. We further present experimental results on a range of datasets illustrating the benefits of our approach, and compare it against alternative solutions: Count-Min sketch and its optimized version called Max-Min sketch, and, on the other hand, Minimal Perfect Hashing. Note, finally, that Set-Min sketch is a general data structure in that it can be used to efficiently represent a mapping of *k*-mers to any type of labels, provided that the number of possible labels is relatively small.

### Applications

Set-Min sketch can have different use cases. In this paper, we explore the probabilistic compression of genomic *k*-mer counter tables in case of large *k* values. Thanks to the characteristic skewed distribution of the frequencies for large *k*s, a Set-Min sketch is able to provide a more space-efficient map than other representations with low errors.

One application of Set-Min sketches, not explored here, is to act as a temporary representation while building more complex structures based on counters. Consider, for example, the exact computation of weighted pairwise distances between all pairs of genomes in a given set. Examples of such distances are the Bray-Curtis similarity measure, see e.g. [12], or Weighted Jaccard similarity estimation [13]. The most naive algorithm is to first produce count tables for each dataset and then compare them pair-wisely to produce the desired output. Instead of storing whole tables, one can store multiple Set-Min sketches together with a presence-absence data structure, such as a Bloom filter. By doing so, the weighted comparison computation is reduced to a single pass through the presence-absence data structure with the counters of a given *k*-mer retrieved on-demand from the Set-Min sketches of the datasets in which the *k*-mer is found.

Another possible application is sharing counter information between different computational units in a distributed setting, where a server oversees multiple less powerful machines. All nodes have access to the same genomic representation (say, a set of contigs), but only the server can efficiently perform *k*-mer counting, while the smaller machines have the task of processing incoming data based on the counts. In this case, the server could send a Set-Min sketch to all its subordinates.

One further feature of Set-Min sketch is its mergeability from redundant maps. A large map can be split into *m* sub-maps without the restriction of having disjoint sets of keys. Even if some maps have redundant information, i.e. share common (key,value) pairs, the Set-Min sketch built by cell-wise union of the *m* sketches will be equivalent to the sketch built from the whole original map. Count-Min sketches do not have this property but instead they are mergeable when constituent maps should be “added up”. In case of redundancy, Count-Min will simply count each repeated item multiple times. In this respect, Set-Min and Count-Min sketches may have complementary uses.

### Outline of the paper

In Section 2, we start by formally introducing the relevant terminology and underlying methods. Section 3 presents the idea and algorithmic foundations of our method. Our results are presented in Section 4 while Section 5 discuss the limitations of our method and possible workarounds. We conclude in Section 6 with a summary of the work.

## 2 Background

### 2.1 k-mer spectrum

A *k*-mer spectrum is a distribution of *k*-mer frequencies across all *k*-mers occurring in the data, showing how many *k*-mers support each frequency value. For large values of *k, k*-mer spectra follow a power-law distribution [10, 11] characterized by a linear-like dependence when represented in the log-log scale. That is, the *k*-mer frequency distribution fits a dependence *f* (*t*) ≈*c·t*^−*a*^, where *a* is usually greater than 2. An example of the spectrum for the human reference genome with *k* equal to 32 is given in Figure 1. According to this distribution, a very large fraction of *k*-mers have very low frequencies, while a few *k*-mers have “unexpectedly” large frequencies. Large values of *a* imply that there are relatively few distinct frequency values with non-zero support, whose number is given by the 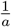 - power of the total number of *k*-mers.

**Figure 1:**
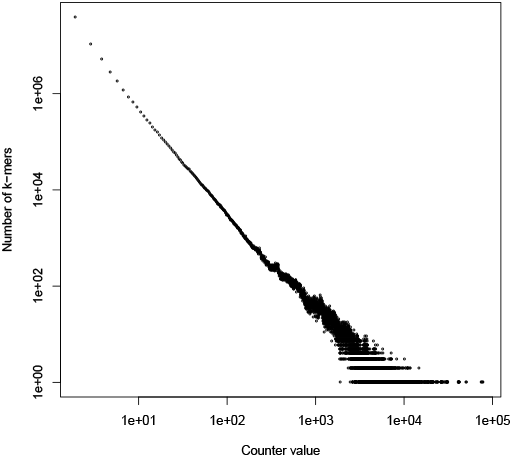
*k*-mer spectrum of the human genome for *k* = 32 in the log-log scale.

### 2.2 Count-Min sketch

Count-Min sketch [14] is a method to compactly represent an associative array ***a*** of counters in an approximated way. Count-Min is especially suitable for the streaming framework, when counters associated to keys can be updated dynamically. A Count-Min sketch is an *R* ×*B* matrix *A* of counters where each row *i* is associated with a hash function *h*_*i*_(·). To store counters and support their dynamic updates given in a stream, Count-Min works as follows. To process an update which is a (key,value) pair (*p, ℓ*), we perform *A*(*i, h*_*i*_(*p*)) = *A*(*i, h*_*i*_(*p*)) + *ℓ* for each row *i*. The (approximate) current counter associated with a key *p* is retrieved as **â** (*p*) = min_*i*_*{h*_*i*_(*p*)*}*.

It has been shown that for a Count-Min sketch built on an associative array (vector) ***a***, with 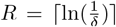 and 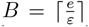 for any given *ε* and *δ*, the over-estimate error of an individual counter is bounded by *ε* ||***a*** ||_1_ with probability at least 1 − *δ*, where ‖;***a*** ‖; _1_ is the *L*_1_-norm of ***a*** [14]. If counts follow a Zipfian rank-frequency distribution with parameter *b >* 1, *B* can be reduced to *O*(*ε*^−1*/b*^) to guarantee the same bounds [15]. If counts follow a power-low distribution with parameter *α >* 1, *B* can be reduced to *O*(*ε*^−1*/α*^) to guarantee the same bounds [15]. In the experimental part of this work (Section 4), we show that these error bounds may not be acceptable for *k*-mer counting applications. Note that *b >* 1 corresponds to 1 < *a* < 2 in the corresponding spectrum distribution [16], while *k*-mer spectra often fit a distribution with *a* ≥2.

Count-Min sketch supports negative updates, i.e. allows *ℓ* < 0 in updates (*p, ℓ*), provided that the cumulative value for each key stays positive. If updates are only positive, there exists a modification of Count-Min leading to a better accuracy, mentioned in [17] (therein attributed to [18]) as *conservative update*. Under this modification, updates for each row *i* are made according to *A*(*i, h*_*i*_(*p*)) = max *{A*(*i, h*_*i*_(*p*)), *â* (*p*) + *ℓ}*, where *â* (*p*) is the current Count-Min estimate of ***a***(*p*). It is easily seen that under this scheme, *â* (*p*) can still only over-estimate ***a***(*p*), but cannot be larger than *â* (*p*) computed by the original Count-Min.

In this paper, we will deal with the static case when the value of any key is given once and never changes after that. In this framework, the conservative update strategy can be further modified by defining updates as *A*(*i, h*_*i*_(*p*)) = max {*A*(*i, h*_*i*_(*p*)), *ℓ*}. We call this variant of the sketch Max-Min. In the static case, Max-Min sketch improves Count-Min without any computational overhead: it simply replaces addition by max in the update rule. As with the conservative update, one can check that estimates by Max-Min can only be over-estimates which, however, don’t exceed estimates by original Count-Min.

### 2.3 Minimal Perfect Hashing

Minimal Perfect Hash Functions (MPHFs) are bijective functions between keys of a set *S* and integers in the range [0, |*S* |−1]. By using hash values as indexes for an external array, it is possible to associate any type of information to the *k*-mers.

The construction of MPHFs can be hyper-graph peeling-based [19, 20] or array-based [21]. The first family of algorithms leads to smaller MPHFs, close to theoretical space lower-bound of 1.44 bits per key, while array-based MPHFs are conceptually simpler and have practical implementations for *k*-mer sets, such as BBHash [22].

## 3 Methods

### 3.1 Set-Min sketch in a nutshell

Assume we are given a set *K* of keys with associated values taken from a set *L* with | *L* | ≪| *K* |. In our case, *K* is a set of *k*-mers and *L* the set of their frequencies, although our method will hold for any set of labels *L*, not necessarily numerical. We want to compactly implement the associative map of (key,value) pairs. A Set-Min sketch is an *R* ×*B* matrix *M* where each bucket is treated as a set, initially empty. Similar to Count-Min sketch, rows in the matrix correspond to hash functions *h*_*i*_, 0 ≤ *i* ≤ *R* − 1, that we assume pairwise independent. At construction time, the key of each (key,value) pair (*p, ℓ*) is hashed by the hash functions to retrieve its buckets, and the value *ℓ* is inserted into each set. Formally, we update *M* (*i, h*_*i*_(*p*)) = *M* (*i, h*_*i*_(*p*)) ∪*{ℓ}*for each row *i*. To retrieve the value associated with a key *p*, we compute the intersection of the corresponding sets, that is ∩_0_≤_*i*_≤_*R* −1_*h*_*i*_(*p*). If the intersection is a singleton, the value is returned. If the intersection is empty, *p* is not present in the map. If the intersection contains more than one value, we have a collision.

Figure 2 shows an example of a Set-Min sketch.

**Figure 2:**
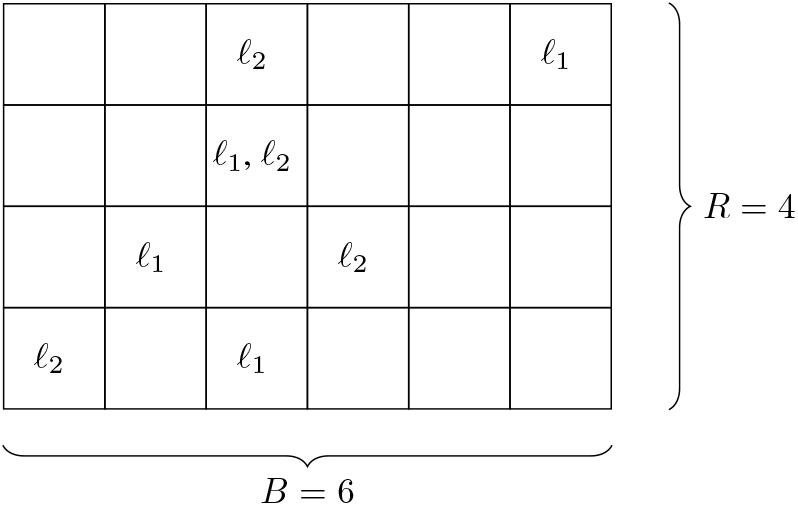
Example of a Set-Min sketch with *L* =*{ℓ*_1_, *ℓ*_2_*}*. Two pairs (*e, ℓ*_1_) and (*f, ℓ*_2_) with *e* ≠ *f* have been inserted into the sketch, with *e, f* hashed to the same bucket at line 2.

**Figure 3:**
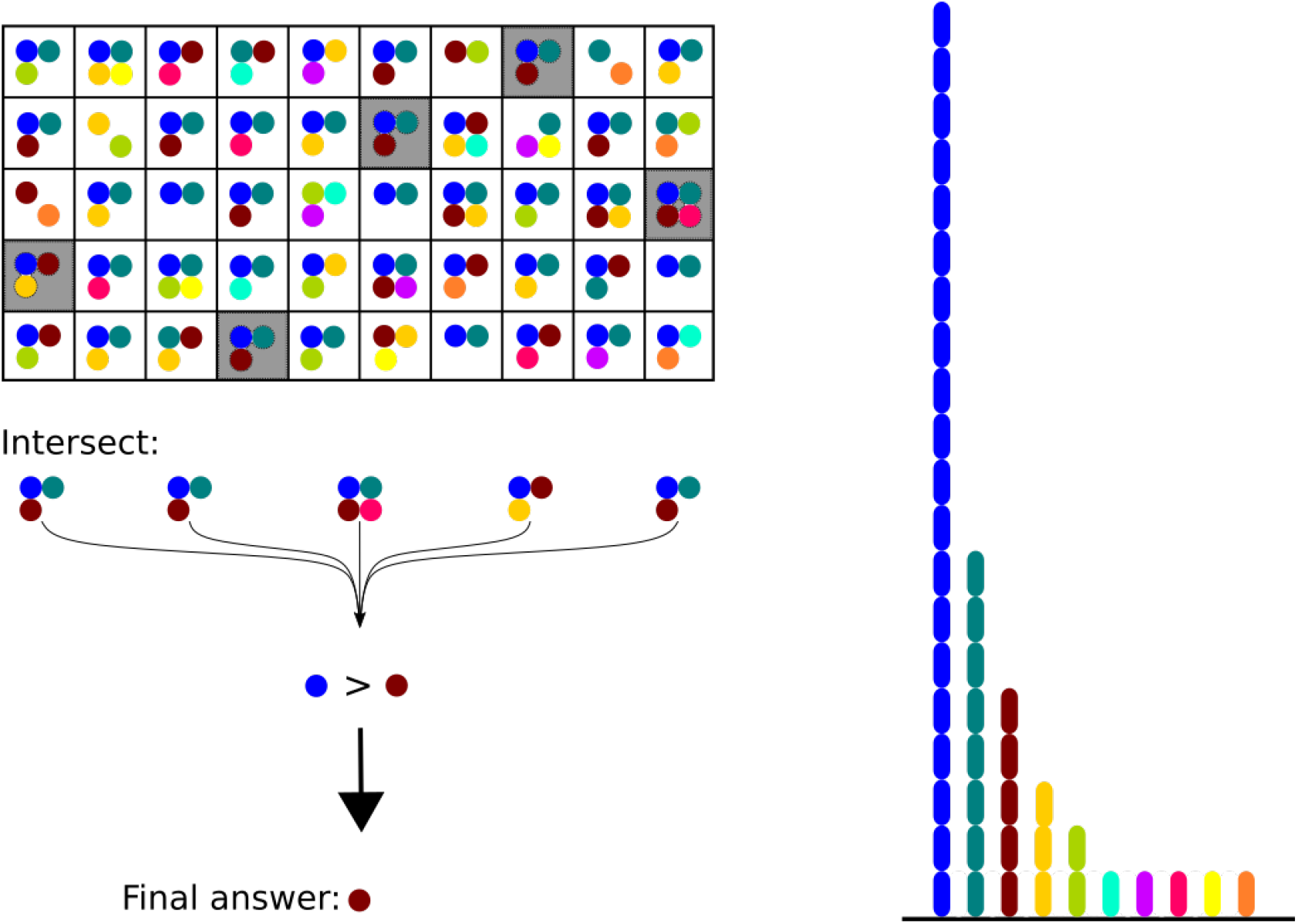
Example of collision resolution in case of multiple items occurring in the intersection. The brown label is returned because it is more rare compared to the blue one.

### 3.2 Dealing with collisions

When we have a collision the choice is guided by the number of *k*-mers supporting each label of the intersection: the label with the smallest support is returned. The rationale for this is that the label with the smallest support has the smallest probability to appear “by chance”, as labels with larger support belong to more buckets in the sketch and are therefore more likely to occur in the intersection by chance. Thus, the algorithm compares spectrum values for all labels in the intersection, and returns the label with the smallest value (ties are broken randomly). If the spectrum is monotonically decreasing (as it is usually the case for large *k*, see Figure 1), then the label returned is simply the largest one among those in the intersection.

In this work we assume that only *k*-mers present in the dataset can be queried. In this case, a query can no longer result in an empty intersection. We further optimize by not storing in the sketch the label *ℓ*_1_ with the largest support. For large *k, ℓ*_1_ is usually 1, which is the frequency of the largest fraction of *k*-mers. *ℓ*_1_ is retrieved implicitly: when the intersection is empty, *ℓ*_1_ is returned. This optimization allows us to save space and will be further discussed later. Note that, with these modification, an error may occur even if the resulting intersection is a singleton but the right label is actually *ℓ*_1_ and not the label obtained.

#### 3.2.1 Bounding the total error

We now show that with Set-Min sketch, we can bound the *total* absolute error over all *k*-mers present in a dataset. Consider a sketch *S* built on a map assigning to each *k*-mer *p* ∈ *K* a value (label) *ℓ*_*p*_ ∈ *L* which is the frequency of *p* in the dataset. We denote by *c*_*£*_ the number of *k*-mers with frequency *ℓ* ∈ *L* (spectrum value).

Consider

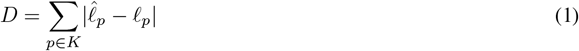

where 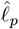 is the label of *p* returned by the sketch. Our goal is to dimension *R* and *B* such that *D* ≤ *ε* ‖ ***a*** ‖ _1_, where *‖* ***a*** *‖* _1_ is the total number of *k*-mers in the dataset (roughly, the dataset size) and 0 < *ϵ* ≤ 1.

Querying *p* returns an incorrect frequency *m* ≠ *ℓ*_*p*_ iff *m* occurs in the intersection and 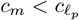. The probability of this event is

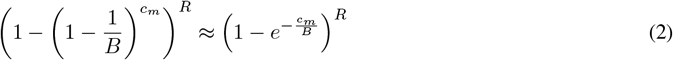

and the expectation of the error when querying *p* is then

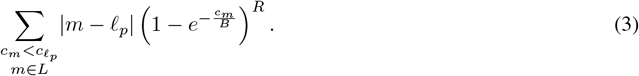

Summing up over all *k*-mers, we obtain

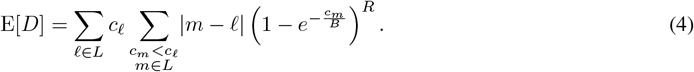

The total number of *k*-mers is | | **a** | | 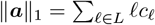. Given 0 < *ε* ≤ 1, our goal is to choose *B* and *R* in order to ensure

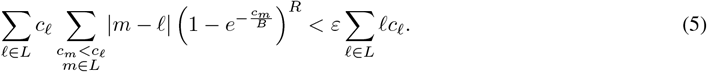

For sufficiently large *k*, the spectrum is monotonically decreasing, i.e. *c*_*m*_ < *c*_*£*_ iff *m > ℓ*. (5) then rewrites to

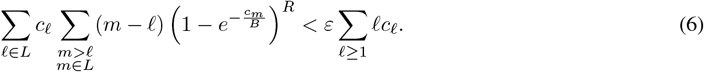

Assume now that the spectrum follows a power-law with large exponent, that is, *c*_*£*_ = *C*·*ℓ*^−*a*^ for some *a >* 2. Note that under this assumption, the number of unique *k*-mers is *c*_1_ = *C*, and the number of all *k*-mers is

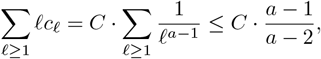

since 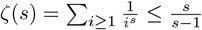 for *s >* 1.

We then have the following result.

##### Theorem 1.

*Given* 0 < *ε* ≤ 1, *if B > C and R, B satisfy*

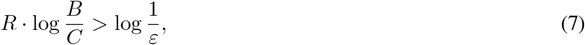

*then (6) holds*.

The proof of Theorem 1 is given in the Appendix. The theorem allows us to dimension the Set-Min sketch. For example, one can set *B* = *αC* for some constant *α >* 1 and 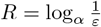.

### 3.3 Computing tighter sketch dimensions

Theorem 1 provides a way to dimension a Set-Min sketch, provided that the spectrum follows a power-law distribution with a sufficiently large parameter *a*. In order to validate these estimates experimentally, and, at the same time, obtain a tool for computing tighter values *B* and *R* for arbitrary spectra, we implemented a simple heuristic hill climbing algorithm to compute those values by directly solving equation 5.

The algorithm, given below, starts with *R* = 1 and some initial value of *B* and then iteratively increments *R* and recomputes (4) until equation (5) holds true. In the implementation, *B* is initially set to 1.44 ×*c*_*max*_, where *c*_*max*_ is the largest spectrum value. After such a value of *R* is found, the algorithm starts decrementing *R* while incrementing *B* to maintain the total space *RB* constant as long as (5) holds. The rationale for this step is to have the smallest possible *R* to reduce query times. Note that, for both loops, there is exist a value of *R* for which the exit condition is satisfied.

#### Algorithm 1

Heuristic to compute *R* and *B*

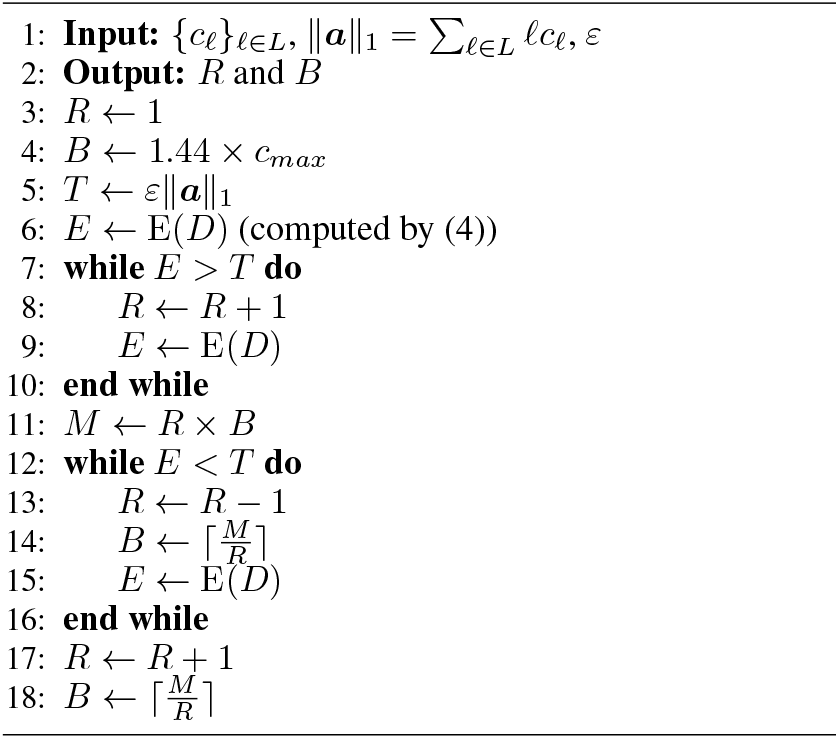

## 4 Results

### 4.1 Implementation

We implemented Set-Min in a software tool named fress, available at https://github.com/yhhshb/fress. The `fress` pipeline compares Set-Min with Count-Min and MPHF implementations. BBHash [22] (https://github.com/rizkg/BBHash) is the only external library required for comparison as Count-Min is implemented within `fress.`

Rather than storing a set in each bucket of the sketch, `fress` only stores an index to an array of involved sets. Note that its purpose is to only validate the approach, and it does not include any complex optimisation, such as multi-threading or bit-packing of the final matrix. Sorted spectra and lists of involved sets are explicitly stored in text format in order to be human-readable and allow for an easier analysis of the results.

### 4.2 Data

We tested Set-Min on six data sets of different size and complexity. Four of them are fully assembled genomes:

- “Sakai” strain of *Escherichia Coli* taken from [23] (NCBI accession number B000007),
- genome of *Drosophila melanogaster* from FlyBase^2^,
- genome of *Gossypium Raimondii* [24] downloadable from AFproject[25],
- human reference genome assembly *GRCh38*^3^.

The other two contain unassembled reads:

- Sakai strain at 5x coverage from AFproject,
- low-coverage human data SRR622461 from the 1000 Genomes Project^4^.

Table 1 summarizes the characteristics of each data set for each value of *k* in our analysis. Observe that, while the number of distinct *k*-mers is comparable to the total number of *k*-mers (data size), the number of distinct *k*-mer counts is small. This is in accordance with the power-low distribution discussed in Section 2.1.

**Table 1:** Data-sheet for the datasets used in our study. Columns Total and Distinct *k*-mers report the number of all and distinct *k*-mers respectively. The Distinct counts column reports the number of distinct *k*-mer frequencies.

| Type | Name | Total $k$-mers [M] | $k$ | Distinct $k$-mers [M] | Distinct counts |
| --- | --- | --- | --- | --- | --- |
| assembled | Sakai | 5.50 | 11 | 2.38 | 69 |
|  |  |  | 15 | 5.23 | 42 |
|  |  |  | 21 | 5.30 | 28 |
|  | melanogaster | 143 | 15 | 101 | 883 |
|  |  |  | 21 | 122 | 710 |
|  |  |  | 27 | 124 | 605 |
|  |  |  | 32 | 125 | 522 |
|  | raimondii | 727 | 15 | 251 | 1944 |
|  |  |  | 21 | 546 | 1019 |
|  |  |  | 27 | 604 | 699 |
|  |  |  | 32 | 632 | 540 |
|  | GRCh38 | 2935 | 15 | 547 | 12718 |
|  |  |  | 21 | 2327 | 10038 |
|  |  |  | 27 | 2483 | 7946 |
|  |  |  | 32 | 2567 | 6651 |
| unassembled | USakai | 25 | 11 | 3.06 | 239 |
|  |  |  | 15 | 9.74 | 143 |
|  |  |  | 21 | 10.0 | 97 |
|  | SRR622461_1 | 7500 | 15 | 676 | 22323 |
|  |  |  | 21 | 3635 | 17211 |
|  |  |  | 27 | 3734 | 13157 |
|  |  |  | 32 | 3703 | 10643 |

### 4.3 Set-Min vs Count-Min sketch

Table 2 compares Set-Min sketch to Count-Min sketch. Dimensions *R* and *B* were computed using Algorithm 1 to insure bound (5) to hold for *ε* = 0.01. Dimensions of Count-Min sketch were set to be the same. To make the comparison fair, the count with the greatest number of k-mers was not inserted into Count-Min sketch, similarly to Set-Min. Zero values are thus interpreted as the non-inserted count.

**Table 2:** Set-Min compared to Count-Min. *R* and *B* are dimensions of Set-Min and Count-Min matrices. *T* is the reference upper bound on the sum of errors equal to *ε* ‖ ***a*** ‖ _1_ (right-hand side of (5)). *E*_*s*_ and *E*_*c*_ are the sum of errors for Set-Min and Count-Min respectively. *N*_*s*_ and *N*_*c*_ are the percentages (rounded to integers) of the fractions of distinct *k*-mers producing an error, for Set-Min and Count-Min respectively. *A*_*s*_ and *A*_*c*_ are respective average errors, with average taken over the number of distinct *k*-mers resulting in an error in the respective sketch.

| *Name* | *k* | *R* | *B* | *T* | *E<sub>s</sub>* | *E<sub>c</sub>* | *N<sub>s</sub>* | *N<sub>c</sub>* | *A<sub>s</sub>* | *A<sub>c</sub>* |
| --- | --- | --- | --- | --- | --- | --- | --- | --- | --- | --- |
| Sakai | 11 | 4 | $1.04 \cdot 10^6$ | $5.50 \cdot 10^4$ | $5.00 \cdot 10^4$ | $1.39 \cdot 10^6$ | 1.8 | 26 | 1.15 | 2.24 |
| Sakai | 15 | 5 | $2.13 \cdot 10^5$ | $5.50 \cdot 10^4$ | $4.66 \cdot 10^4$ | $2.38 \cdot 10^5$ | 0.9 | 4 | 1.01 | 1.1 |
| Sakai | 21 | 5 | $1.20 \cdot 10^5$ | $5.50 \cdot 10^4$ | $5.29 \cdot 10^4$ | $4.57 \cdot 10^5$ | 0.9 | 7 | 1.05 | 1.18 |
| USakai | 11 | 5 | $4.58 \cdot 10^5$ | $2.57 \cdot 10^5$ | $1.99 \cdot 10^5$ | $7.44 \cdot 10^7$ | 2.6 | 99 | 2.46 | 24.6 |
| USakai | 15 | 4 | $4.79 \cdot 10^6$ | $2.49 \cdot 10^5$ | $2.45 \cdot 10^5$ | $8.01 \cdot 10^6$ | 1.9 | 33 | 1.31 | 2.53 |
| USakai | 21 | 5 | $3.99 \cdot 10^6$ | $2.38 \cdot 10^5$ | $2.00 \cdot 10^5$ | $7.91 \cdot 10^6$ | 1.6 | 34 | 1.21 | 2.35 |
| melanogaster | 15 | 5 | $1.79 \cdot 10^7$ | $1.43 \cdot 10^6$ | $1.39 \cdot 10^6$ | $8.35 \cdot 10^6$ | 1.4 | 7 | 1 | 1.25 |
| melanogaster | 21 | 4 | $4.69 \cdot 10^6$ | $1.43 \cdot 10^6$ | $1.41 \cdot 10^6$ | $2.26 \cdot 10^7$ | 1.1 | 12 | 1.03 | 1.61 |
| melanogaster | 27 | 5 | $3.72 \cdot 10^6$ | $1.43 \cdot 10^6$ | $1.11 \cdot 10^6$ | $2.28 \cdot 10^7$ | 0.9 | 12 | 1.01 | 1.48 |
| melanogaster | 32 | 5 | $3.81 \cdot 10^6$ | $1.42 \cdot 10^6$ | $1.11 \cdot 10^6$ | $2.10 \cdot 10^7$ | 0.9 | 12 | 1.01 | 1.44 |
| raimondii | 15 | 4 | $8.36 \cdot 10^7$ | $7.27 \cdot 10^6$ | $5.65 \cdot 10^6$ | $1.50 \cdot 10^8$ | 2.1 | 27 | 1.08 | 2.25 |
| raimondii | 21 | 4 | $8.47 \cdot 10^7$ | $7.27 \cdot 10^6$ | $5.73 \cdot 10^6$ | $4.09 \cdot 10^7$ | 1 | 6 | 1.01 | 1.3 |
| raimondii | 27 | 4 | $7.14 \cdot 10^7$ | $7.27 \cdot 10^6$ | $6.49 \cdot 10^6$ | $3.56 \cdot 10^7$ | 1.1 | 5 | 1.01 | 1.22 |
| raimondii | 32 | 4 | $6.38 \cdot 10^7$ | $7.27 \cdot 10^6$ | $6.87 \cdot 10^6$ | $3.15 \cdot 10^7$ | 1.1 | 4 | 1.01 | 1.17 |
| GRCh38 | 15 | 3 | $2.26 \cdot 10^8$ | $2.93 \cdot 10^7$ | $2.78 \cdot 10^7$ | $1.36 \cdot 10^9$ | 2.9 | 52 | 1.78 | 4.74 |
| GRCh38 | 21 | 4 | $1.42 \cdot 10^8$ | $2.93 \cdot 10^7$ | $2.60 \cdot 10^7$ | $1.65 \cdot 10^8$ | 1.1 | 6 | 1.01 | 1.23 |
| GRCh38 | 27 | 4 | $1.07 \cdot 10^8$ | $2.93 \cdot 10^7$ | $2.82 \cdot 10^7$ | $1.91 \cdot 10^8$ | 1.1 | 6 | 1.01 | 1.25 |
| GRCh38 | 32 | 4 | $9.84 \cdot 10^7$ | $2.93 \cdot 10^7$ | $2.92 \cdot 10^7$ | $1.85 \cdot 10^8$ | 1.1 | 6 | 1.01 | 1.22 |
| SRR622461 | 15 | 4 | $1.17 \cdot 10^8$ | $7.90 \cdot 10^7$ | $4.93 \cdot 10^7$ | $1.34 \cdot 10^{10}$ | 3.3 | 96 | 2.2 | 20.68 |
| SRR622461 | 21 | 3 | $2.39 \cdot 10^9$ | $7.35 \cdot 10^7$ | $7.09 \cdot 10^7$ | $5.63 \cdot 10^8$ | 1.8 | 9 | 1.11 | 1.68 |
| SRR622461 | 27 | 4 | $1.78 \cdot 10^9$ | $6.80 \cdot 10^7$ | $4.92 \cdot 10^7$ | $4.58 \cdot 10^8$ | 1.3 | 8 | 1.04 | 1.53 |
| SRR622461 | 32 | 4 | $1.72 \cdot 10^9$ | $6.34 \cdot 10^7$ | $4.95 \cdot 10^7$ | $4.01 \cdot 10^8$ | 1.3 | 7 | 1.03 | 1.48 |

For ease of comparison, column *T* reports the threshold *ε* ‖ ***a*** ‖ _1_ given to Algorithm 1. Columns *E*_*s*_ and *E*_*c*_ report the actual total sum of errors for Set-Min and Count-Min, respectively. In all reported cases *E*_*s*_ < *T*, as expected. The total error of Count-Min, *E*_*c*_ is, most of the time, one order of magnitude larger than *E*_*s*_. For SRR622461 with *k* = 15, it even exceeds the total number of *k*-mers ‖ ***a*** ‖ _1_ in the dataset.

The average error of Set-Min is, in most cases, very close to 1, which suggests that the overwhelming majority of collisions occur between successive counts such as 1 and 2 – the most abundant ones in the spectra considered here. The average error of Count-Min is bigger but of the same order of magnitude, except for small *k* and unassembled datasets.

On the other hand, the fraction of *k*-mers producing an error is in striking contrast: in case of Set-Min, about only 1-3% of distinct *k*-mers produce an error, while for Count-Min, this fraction is much larger. This shows that Count-Min cannot be used when most of *k*-mer counts are expected to be retrieved precisely, for comparable sketch sizes.

### 4.4 Set-Min vs Max-Min sketch

We compared Set-Min to Max-Min – an optimized version of Count-Min (see Sect. **??**). Note that in the case when the *k*-mer spectrum is strictly decreasing for increasing *k*-mer counts, the maximum count corresponds to the one with the smallest support. In the general case, we use a variant when, instead of using the ordinary order, counts are ordered according to the support size. That is, updates are performed by keeping the label with the minimum number of *k*-mers in the *k*-mer spectrum, and a query returns the label with the smallest such number.

Table 3 compares Set-Min with Max-Min. As expected from theoretical considerations, the performance of Max-Min in terms of the average error and the sum of errors is better than for regular Count-Min, but worse than for Set-Min. The same behaviour is observed for the number of *k*-mers having an erroneously estimated frequency. Therefore, Max-Min falls in-between Set-Min and Count-Min, providing a simple and inexpensive practical method to enhance the latter, without reaching the accuracy of the former. Altogether, Table 3 shows that the performance of Max-Min is closer to Count-Min than to Set-Min.

**Table 3:** Set-Min compared to Max-Min sketch. Columns *E*_*c*_, *N*_*c*_, *A*_*c*_ are replaced by *E*_*m*_, *N*_*m*_, *A*_*m*_ with the same meaning as their Table 2 counterparts.

| *Name* | *k* | *R* | *B* | *T* | $E_s$ | $E_m$ | $N_s$ | $N_m$ | $A_s$ | $A_m$ |
| --- | --- | --- | --- | --- | --- | --- | --- | --- | --- | --- |
| Sakai | 11 | 4 | $1.04 \cdot 10^6$ | $5.50 \cdot 10^4$ | $5.00 \cdot 10^4$ | $4.15 \cdot 10^5$ | 1.8 | 13.5 | 1.15 | 1.29 |
| Sakai | 15 | 5 | $2.13 \cdot 10^5$ | $5.50 \cdot 10^4$ | $4.66 \cdot 10^4$ | $2.13 \cdot 10^5$ | 0.9 | 4 | 1.01 | 1.01 |
| Sakai | 21 | 5 | $1.20 \cdot 10^5$ | $5.50 \cdot 10^4$ | $5.29 \cdot 10^4$ | $4.02 \cdot 10^5$ | 0.9 | 7.2 | 1.05 | 1.06 |
| USakai | 11 | 5 | $4.58 \cdot 10^5$ | $2.57 \cdot 10^5$ | $1.99 \cdot 10^5$ | $1.24 \cdot 10^7$ | 2.6 | 63.5 | 2.46 | 6.4 |
| USakai | 15 | 4 | $4.79 \cdot 10^6$ | $2.49 \cdot 10^5$ | $2.45 \cdot 10^5$ | $1.72 \cdot 10^6$ | 1.9 | 13.3 | 1.31 | 1.33 |
| USakai | 21 | 5 | $3.99 \cdot 10^6$ | $2.38 \cdot 10^5$ | $2.00 \cdot 10^5$ | $1.76 \cdot 10^6$ | 1.6 | 14.2 | 1.21 | 1.24 |
| melanogaster | 15 | 5 | $1.79 \cdot 10^7$ | $1.43 \cdot 10^6$ | $1.39 \cdot 10^6$ | $5.74 \cdot 10^6$ | 1.4 | 5.6 | 1 | 1.02 |
| melanogaster | 21 | 4 | $4.69 \cdot 10^6$ | $1.43 \cdot 10^6$ | $1.41 \cdot 10^6$ | $1.78 \cdot 10^7$ | 1.1 | 11.2 | 1.03 | 1.31 |
| melanogaster | 27 | 5 | $3.72 \cdot 10^6$ | $1.43 \cdot 10^6$ | $1.11 \cdot 10^6$ | $1.78 \cdot 10^7$ | 0.9 | 12.1 | 1.01 | 1.2 |
| melanogaster | 32 | 5 | $3.81 \cdot 10^6$ | $1.42 \cdot 10^6$ | $1.11 \cdot 10^6$ | $1.65 \cdot 10^7$ | 0.9 | 11.3 | 1.01 | 1.17 |
| raimondii | 15 | 4 | $8.36 \cdot 10^7$ | $7.27 \cdot 10^6$ | $5.65 \cdot 10^6$ | $5.81 \cdot 10^7$ | 2.1 | 16.8 | 1.08 | 1.38 |
| raimondii | 21 | 4 | $8.47 \cdot 10^7$ | $7.27 \cdot 10^6$ | $5.73 \cdot 10^6$ | $3.05 \cdot 10^7$ | 1 | 5.2 | 1.01 | 1.07 |
| raimondii | 27 | 4 | $7.14 \cdot 10^7$ | $7.27 \cdot 10^6$ | $6.49 \cdot 10^6$ | $2.84 \cdot 10^7$ | 1.1 | 4.5 | 1.01 | 1.05 |
| raimondii | 32 | 4 | $6.38 \cdot 10^7$ | $7.27 \cdot 10^6$ | $6.87 \cdot 10^6$ | $2.61 \cdot 10^7$ | 1.1 | 4 | 1.01 | 1.03 |
| GRCh38 | 15 | 3 | $2.26 \cdot 10^8$ | $2.93 \cdot 10^7$ | $2.78 \cdot 10^7$ | $4.07 \cdot 10^8$ | 2.9 | 27.8 | 1.78 | 2.67 |
| GRCh38 | 21 | 4 | $1.42 \cdot 10^8$ | $2.93 \cdot 10^7$ | $2.60 \cdot 10^7$ | $1.38 \cdot 10^8$ | 1.1 | 5.5 | 1.01 | 1.08 |
| GRCh38 | 27 | 4 | $1.07 \cdot 10^8$ | $2.93 \cdot 10^7$ | $2.82 \cdot 10^7$ | $1.62 \cdot 10^8$ | 1.1 | 6 | 1.01 | 1.09 |
| GRCh38 | 32 | 4 | $9.84 \cdot 10^7$ | $2.93 \cdot 10^7$ | $2.92 \cdot 10^7$ | $1.58 \cdot 10^8$ | 1.1 | 5.7 | 1.01 | 1.08 |
| SRR622461 | 15 | 4 | $1.17 \cdot 10^8$ | $7.90 \cdot 10^7$ | $4.93 \cdot 10^7$ | $3.31 \cdot 10^9$ | 3.3 | 61.8 | 2.2 | 7.92 |
| SRR622461 | 21 | 3 | $2.39 \cdot 10^9$ | $7.35 \cdot 10^7$ | $7.09 \cdot 10^7$ | $2.42 \cdot 10^8$ | 1.8 | 5.9 | 1.11 | 1.13 |
| SRR622461 | 27 | 4 | $1.78 \cdot 10^9$ | $6.80 \cdot 10^7$ | $4.92 \cdot 10^7$ | $2.05 \cdot 10^8$ | 1.3 | 5.2 | 1.04 | 1.06 |
| SRR622461 | 32 | 4 | $1.72 \cdot 10^9$ | $6.34 \cdot 10^7$ | $4.95 \cdot 10^7$ | $1.89 \cdot 10^8$ | 1.3 | 4.9 | 1.03 | 1.04 |

This is because, by keeping the maximum element in each bucket, we are reducing each set of Set-Min to a single element opening the possibility of increased collisions by potentially sharing a given maximum element between unrelated *k*-mers. The intersection operation performed by Set-Min during query is thus strictly necessary, to guarantee the desired error bounds.

### 4.5 Set-Min sketch vs KMC output

Not surprisingly, Set-Min achieves better memory consumptions than KMC in all our tests (columns *M*_*kmc*_ and *M*_*s*_ of Table 4). Values of *R* and *B* do not change from Table 2. The compression rate is variable: from a small factor to two orders of magnitude. The best compression is achieved for larger values of *k* and assembled genomes. The former is primarily explained by the decreasing number of distinct counts, due to the power-law behaviour. As for the difference between assembled genomes and sequencing data, we will discuss it in more details in Section 5.

**Table 4:** Set-Min (*ε* = 0.01) compared to KMC and BBHash (*γ* = 1). All memory is reported in bytes. Column *M*_*kmc*_, *M*_*s*_, *M*_*bball*_ are the memory requirements to have a fully functional map between *k*-mers and their frequencies when applying KMC, Set-Min sketch and BBHash. *M*_*bbhash*_ is the memory of the hash function produced by BBHash without the external array of frequencies, *C*_*s*_ and *C*_*bball*_ are the compressed versions of *M*_*s*_ and *M*_*bball*_ using gzip.

| *Name* | *k* | $M_{kmc}$ | $M_s$ | $C_s$ | $M_{bbhash}$ | $M_{bball}$ | $C_{bball}$ |
| --- | --- | --- | --- | --- | --- | --- | --- |
| Sakai | 11 | $1.21 \cdot 10^7$ | $5.75 \cdot 10^6$ | $3.41 \cdot 10^6$ | $9.13 \cdot 10^5$ | $3.00 \cdot 10^6$ | $2.01 \cdot 10^6$ |
| Sakai | 15 | $3.80 \cdot 10^7$ | $1.20 \cdot 10^6$ | $5.55 \cdot 10^5$ | $2.06 \cdot 10^6$ | $5.99 \cdot 10^6$ | $2.18 \cdot 10^6$ |
| Sakai | 21 | $4.77 \cdot 10^7$ | $6.77 \cdot 10^5$ | $3.57 \cdot 10^5$ | $2.03 \cdot 10^6$ | $6.00 \cdot 10^6$ | $2.09 \cdot 10^6$ |
| USakai | 11 | $1.54 \cdot 10^7$ | $1.49 \cdot 10^7$ | $7.99 \cdot 10^6$ | $1.17 \cdot 10^6$ | $4.61 \cdot 10^6$ | $3.68 \cdot 10^6$ |
| USakai | 15 | $6.95 \cdot 10^7$ | $2.87 \cdot 10^7$ | $1.54 \cdot 10^7$ | $3.72 \cdot 10^6$ | $1.35 \cdot 10^7$ | $8.34 \cdot 10^6$ |
| USakai | 21 | $8.53 \cdot 10^7$ | $3.00 \cdot 10^7$ | $1.61 \cdot 10^7$ | $3.91 \cdot 10^6$ | $1.39 \cdot 10^7$ | $8.40 \cdot 10^6$ |
| melanogaster | 15 | $7.08 \cdot 10^8$ | $1.68 \cdot 10^8$ | $5.74 \cdot 10^7$ | $3.87 \cdot 10^7$ | $2.15 \cdot 10^8$ | $5.76 \cdot 10^7$ |
| melanogaster | 21 | $9.80 \cdot 10^8$ | $3.55 \cdot 10^7$ | $1.62 \cdot 10^7$ | $4.66 \cdot 10^7$ | $2.45 \cdot 10^8$ | $5.22 \cdot 10^7$ |
| melanogaster | 27 | $1.24 \cdot 10^9$ | $3.75 \cdot 10^7$ | $1.77 \cdot 10^7$ | $4.73 \cdot 10^7$ | $2.48 \cdot 10^8$ | $5.26 \cdot 10^7$ |
| melanogaster | 32 | $1.37 \cdot 10^9$ | $3.84 \cdot 10^7$ | $1.78 \cdot 10^7$ | $4.90 \cdot 10^7$ | $2.36 \cdot 10^8$ | $5.38 \cdot 10^7$ |
| raimondii | 15 | $1.76 \cdot 10^9$ | $7.54 \cdot 10^8$ | $3.34 \cdot 10^8$ | $9.59 \cdot 10^7$ | $5.98 \cdot 10^8$ | $2.11 \cdot 10^8$ |
| raimondii | 21 | $4.37 \cdot 10^9$ | $6.78 \cdot 10^8$ | $2.08 \cdot 10^8$ | $2.16 \cdot 10^8$ | $1.24 \cdot 10^9$ | $2.90 \cdot 10^8$ |
| raimondii | 27 | $6.04 \cdot 10^9$ | $5.36 \cdot 10^8$ | $1.62 \cdot 10^8$ | $2.37 \cdot 10^8$ | $1.29 \cdot 10^9$ | $2.94 \cdot 10^8$ |
| raimondii | 32 | $6.96 \cdot 10^9$ | $4.79 \cdot 10^8$ | $1.36 \cdot 10^8$ | $2.70 \cdot 10^8$ | $1.38 \cdot 10^9$ | $3.10 \cdot 10^8$ |
| GRCh38 | 15 | $3.83 \cdot 10^9$ | $1.70 \cdot 10^9$ | $9.32 \cdot 10^8$ | $2.09 \cdot 10^8$ | $1.65 \cdot 10^9$ | $5.67 \cdot 10^8$ |
| GRCh38 | 21 | $1.86 \cdot 10^{10}$ | $1.28 \cdot 10^9$ | $3.58 \cdot 10^8$ | $9.32 \cdot 10^8$ | $6.46 \cdot 10^9$ | $1.03 \cdot 10^9$ |
| GRCh38 | 27 | $2.48 \cdot 10^{10}$ | $9.66 \cdot 10^8$ | $2.79 \cdot 10^8$ | $9.86 \cdot 10^8$ | $6.57 \cdot 10^9$ | $1.05 \cdot 10^9$ |
| GRCh38 | 32 | $2.82 \cdot 10^{10}$ | $8.38 \cdot 10^8$ | $2.50 \cdot 10^8$ | $1.08 \cdot 10^9$ | $6.54 \cdot 10^9$ | $1.12 \cdot 10^9$ |
| SRR622461 | 15 | $4.73 \cdot 10^9$ | $2.00 \cdot 10^9$ | $1.51 \cdot 10^9$ | $2.59 \cdot 10^8$ | $2.12 \cdot 10^9$ | $8.38 \cdot 10^8$ |
| SRR622461 | 21 | $2.91 \cdot 10^{10}$ | $1.61 \cdot 10^{10}$ | $3.66 \cdot 10^9$ | $1.48 \cdot 10^9$ | $1.10 \cdot 10^{10}$ | $2.71 \cdot 10^9$ |
| SRR622461 | 27 | $3.73 \cdot 10^{10}$ | $1.60 \cdot 10^{10}$ | $3.88 \cdot 10^9$ | $1.48 \cdot 10^9$ | $1.08 \cdot 10^{10}$ | $2.66 \cdot 10^9$ |
| SRR622461 | 32 | $4.07 \cdot 10^{10}$ | $1.46 \cdot 10^{10}$ | $3.59 \cdot 10^9$ | $1.57 \cdot 10^9$ | $1.04 \cdot 10^{10}$ | $2.65 \cdot 10^9$ |

### 4.6 Set-Min sketch vs MPHFs

Table 4 also reports the space usage for `BBHash` with parameter *γ* = 1 to obtain the best memory-optimized hash functions. Column *M*_*bbhash*_ is the space (in bytes) required by the hash function only, while *M*_*bball*_ is the space required by the hash function plus the external array of frequencies.

As in the previous case, Set-Min sketch is more memory-efficient when *k* is large, taking about an order of magnitude less memory than a MPHF. For small values of *k*, `BBHash` takes slightly less space and, being exact, may therefore be the preferable choice. However, one should keep in mind that MPHF does not support updates, while a Set-Min sketch is updatable to a certain extent with new (*k*-mer,count) pairs, and also mergeable with another possibly redundant map. For long-term storage through gzip compression of the sketches (columns *C*_*s*_ and *C*_*bball*_) Set-Min and MPHFs give equivalent results.

The behaviour of the unassembled datasets is of particular interest. Even for large *k*’s, MPHF appears to be a better choice for this type of data. The causes of this phenomenon and possible solutions are discussed in Section 5.

## 5 Discussion

### 5.1 Unassembled datasets

As seen in Table 4, for the unassembled datasets, Set-Min sketch does not seem to have an advantage in memory usage, even for large *k*’s. We found that this is due to low-count *k*-mers, specifically to *k*-mers whose count does not exceed the sequencing coverage. It is known that for Illumina sequencing, sequencing errors produce a linear growth of the number of new distinct *k*-mers (for large *k*) depending on the coverage (see e.g. Figure 2(b) of [26]). Frequencies of these “erroneous” *k*-mers do not have the same statistical behaviour as *bona fide k*-mers, in particular first spectrum values do not decay at the same rate as the rest of the spectrum.

Figure 4 shows spectra of the unassembled USakai and SRR622461 datasets. One can observe a slower decay behaviour for a first few spectrum values. In this situation, Algorithm 1 generates a lot of rows just to make the sketch able to distinguish, with required precision, between small frequency values.

Note that in practice, distinguishing between small frequencies is often irrelevant. For example, many read assemblers simply discard low-frequency reads as a way to de-noise the data. In the case of Set-Min, it is possible to collapse together the first *m* columns of the spectrum by assigning to all *k*-mers in this subset the same frequency. This would considerably reduce the sketch size. Formally, the error guarantee (5) would not hold anymore but most of newly introduced errors would be small (typically, equal to 1) and occur for low counts only.

**Figure 4:**
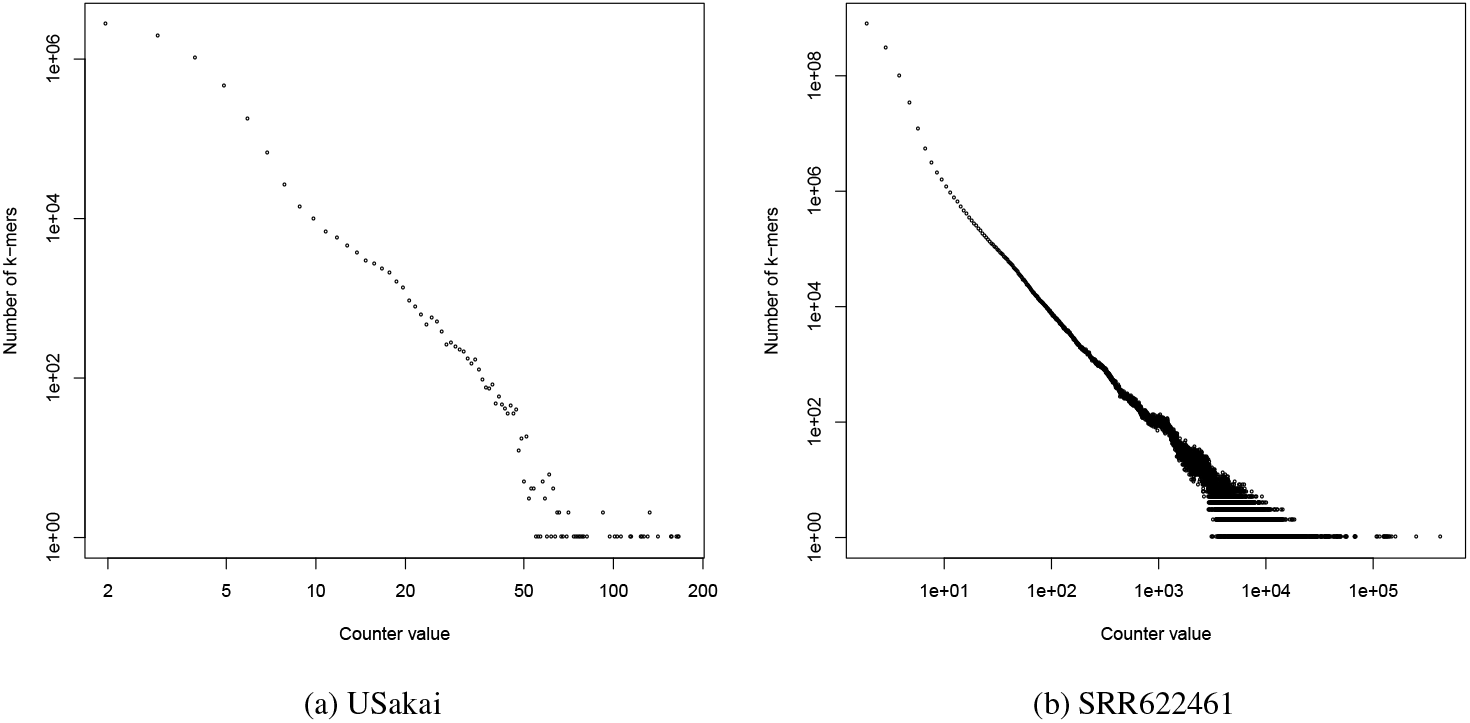
Spectra for the unassembled data sets. Both plots are in log-log scale. *k* = 21 for USakai, *k* = 32 for SRR622461.

To check the above, we constructed a Set-Min sketch for SRR622461 with dimensions (*R, B*) = (4, 3310557), merging together the first five columns of the sorted spectrum and assigning count 5 to all merged *k*-mers. While the final sum of errors was well above the theoretical limit (10^9^ against 7·10^6^), the maximum and average error were respectively 55 and 2.8. In many applications this error level could be acceptable.

### 5.2 Presence-absence information

As introduced in Section 3.2, in this work we assumed that only *k*-mers present in the dataset can be queried. This assumption allowed us to discard the largest value of the spectrum corresponding to unique *k*-mers, thereby saving more space than a complete sketch.

Set-Min sketches can seamlessly work without this assumption, but the space required for storing *k*-mer counters may not be competitive to other solutions. An alternative is to build an additional data structure, specifically optimized to answer presence/absence queries (such as a Bloom filter), as a complement to Set-Min sketches.

Another scenario occurs when working with multiple datasets of very high similarity, such as a large collection of bacterial strains or a collection of RNA-seq data [27, 28]. In this case, it might be beneficial to build a Bloom filter for the *k*-mers present in the union of the datasets, and maintain multiple Set-Min sketches to represent *k*-mer counts in each dataset.

Note that Set-Min sketches can be also used for long-term storage and transmission of the *k*-mer composition of a dataset augmented with count information. The *k*-mers of the dataset can be reassembled into simplitigs [8] with a Set-Min sketch storing the (approximated) frequencies. The full count table can be restored from the simplitigs and the sketch.

## 6 Conclusions

We presented Set-Min sketch – a novel sketching method inspired by the Count-Min sketch. Its primary use is to associate keys to labels without explicitly storing the former. In this paper, we demonstrated the performance of Set-Min sketch for storing *k*-mer counts information, where the distribution of labels (*k*-mer counts) follows a power-law. Under this assumption, we proposed simple bounds for a Set-Min sketch that guarantee the total error sum to be within an *ε* fraction of the total number of *k*-mers in the dataset.

We showed that Set-Min sketch allows us to save space compared to the raw output of the popular KMC *k*-mer counting tool when applied to labels following a skewed distribution, at the price of a very modest error rate. Memory efficiency is particularly significant in case of whole-genome data and large values of *k*, where it can reach reductions of two orders of magnitude. Set-Min has been shown to be more space efficient than the MPHF-based soluton for large values of *k*. For smaller *k*’s and for unassembled datasets, MPHFs remain competitives. In the latter case, it is possible to relax the error guarantee of Set-Min, if low counts are not important to achieve smaller sketch sizes, if needed. Finally, when compared to Count-Min sketches of comparable dimensions, our sketch achieves better point-query errors thanks to the distribution-aware dimensioning performed on the *k*-mer spectrum, and the reliance on already computed count tables as input.

## Acknowledgements

GK was partially funded by RFBR, project 20-07-00652, and joint RFBR and JSPS project 20-51-50007.

## Supplemental Material: Set-Min sketch

**Proof of Theorem 1**

Our goal is to estimate

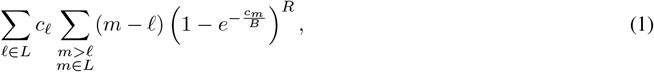

where *c_ℓ_* = *C ℓ*^−*a*^. We assume *B > C* and approximate 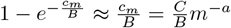. We further lower-approximate (1) by replacing sums by integrals, thus obtaining

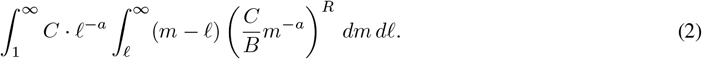

Routine computation of the integral yields

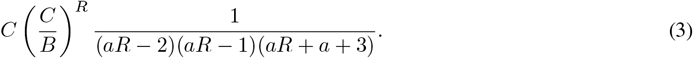

The inequality of the Theorem becomes

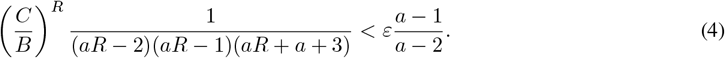

The Theorem follows.

**Performance comparison for epsilon = 0.01**

**Table 1:** Full table comparing the performance of Set-Min sketch constructed for *E* = 0.01 to all other variables for all possible *k*s. *R* = number of rows, *B* = number of columns, *T* = theoretical maximum error bound, *E*_*s*_ = sum of errors for Set-Min, *N*_*s*_ = total number of collisions for Set-Min, *A*_*s*_ = average point-query error (E_S_*/N*_*S*_),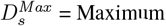 point-query error for Set-Min, *E*_*c*_ = sum of errors for Count-Min, *N*_*c*_ = total number of collisions for Count-Min, *A*_*c*_ = average point-query error (*E*_*c*_*/N*_*c*_), 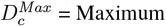 point-query error for Count-Min

| *Name* | *k* | *R* | *B* | *T* | *E<sub>s</sub>* | *N<sub>s</sub>* | *A<sub>s</sub>* | *D<sub>s</sub><sup>Max</sup>* | *E<sub>c</sub>* | *N<sub>c</sub>* | *A<sub>c</sub>* | *D<sub>c</sub><sup>Max</sup>* |
| --- | --- | --- | --- | --- | --- | --- | --- | --- | --- | --- | --- | --- |
| Sakai | 8 | 7 | 1,020 | $5.50 \cdot 10^4$ | 42,059 | $3.00 \cdot 10^0$ | 21.56 | 122 | $2.80 \cdot 10^8$ | $1.00 \cdot 10^2$ | 4,284.98 | 6,129 |
| Sakai | 11 | 4 | 1,042,202 | $5.50 \cdot 10^4$ | 49,953 | $1.80 \cdot 10^0$ | 1.15 | 7 | $1.39 \cdot 10^6$ | $2.60 \cdot 10^1$ | 2.24 | 15 |
| Sakai | 15 | 5 | 212,522 | $5.50 \cdot 10^4$ | 46,619 | $9.00 \cdot 10^{-1}$ | 1.01 | 2 | $2.38 \cdot 10^5$ | $4.00 \cdot 10^0$ | 1.1 | 7 |
| Sakai | 21 | 5 | 120,006 | $5.50 \cdot 10^4$ | 52,852 | $9.00 \cdot 10^{-1}$ | 1.05 | 4 | $4.57 \cdot 10^5$ | $7.00 \cdot 10^0$ | 1.18 | 7 |
| USakai | 8 | 7 | 276 | $2.62 \cdot 10^5$ | $2.30 \cdot 10^5$ | $3.50 \cdot 10^0$ | 101.03 | 623 | $5.55 \cdot 10^9$ | $1.00 \cdot 10^2$ | 84,769.1 | $1.00 \cdot 10^5$ |
| USakai | 11 | 5 | 458,224 | $2.57 \cdot 10^5$ | $1.99 \cdot 10^5$ | $2.60 \cdot 10^0$ | 2.46 | 23 | $7.44 \cdot 10^7$ | $9.90 \cdot 10^1$ | 24.6 | 121 |
| USakai | 15 | 4 | 4,786,091 | $2.49 \cdot 10^5$ | $2.45 \cdot 10^5$ | $1.90 \cdot 10^0$ | 1.31 | 5 | $8.01 \cdot 10^6$ | $3.30 \cdot 10^1$ | 2.53 | 16 |
| USakai | 21 | 5 | 3,992,620 | $2.38 \cdot 10^5$ | $2.00 \cdot 10^5$ | $1.60 \cdot 10^0$ | 1.21 | 4 | $7.91 \cdot 10^6$ | $3.40 \cdot 10^1$ | 2.35 | 13 |
| melanogaster | 15 | 5 | 17,850,639 | $1.43 \cdot 10^6$ | $1.39 \cdot 10^6$ | $1.40 \cdot 10^0$ | 1 | 4 | $8.35 \cdot 10^6$ | $7.00 \cdot 10^0$ | 1.25 | 18 |
| melanogaster | 21 | 4 | 4,694,088 | $1.43 \cdot 10^6$ | $1.41 \cdot 10^6$ | $1.10 \cdot 10^0$ | 1.03 | 13 | $2.26 \cdot 10^7$ | $1.20 \cdot 10^1$ | 1.61 | 55 |
| melanogaster | 27 | 5 | 3,717,540 | $1.43 \cdot 10^6$ | $1.11 \cdot 10^6$ | $9.00 \cdot 10^{-1}$ | 1.01 | 7 | $2.28 \cdot 10^7$ | $1.20 \cdot 10^1$ | 1.48 | 34 |
| melanogaster | 32 | 5 | 3,814,440 | $1.42 \cdot 10^6$ | $1.11 \cdot 10^6$ | $9.00 \cdot 10^{-1}$ | 1.01 | 8 | $2.10 \cdot 10^7$ | $1.20 \cdot 10^1$ | 1.44 | 30 |
| raimondii | 15 | 4 | 83,575,168 | $7.27 \cdot 10^6$ | $5.65 \cdot 10^6$ | $2.10 \cdot 10^0$ | 1.08 | 11 | $1.50 \cdot 10^8$ | $2.70 \cdot 10^1$ | 2.25 | 65 |
| raimondii | 21 | 4 | 84,742,080 | $7.27 \cdot 10^6$ | $5.73 \cdot 10^6$ | $1.00 \cdot 10^0$ | 1.01 | 9 | $4.09 \cdot 10^7$ | $6.00 \cdot 10^0$ | 1.3 | 36 |
| raimondii | 27 | 4 | 71,403,946 | $7.27 \cdot 10^6$ | $6.49 \cdot 10^6$ | $1.10 \cdot 10^0$ | 1.01 | 7 | $3.56 \cdot 10^7$ | $5.00 \cdot 10^0$ | 1.22 | 32 |
| raimondii | 32 | 4 | 63,793,114 | $7.27 \cdot 10^6$ | $6.87 \cdot 10^6$ | $1.10 \cdot 10^0$ | 1.01 | 7 | $3.15 \cdot 10^7$ | $4.00 \cdot 10^0$ | 1.17 | 22 |
| GRCh38 | 15 | 3 | 225,505,316 | $2.93 \cdot 10^7$ | $2.78 \cdot 10^7$ | $2.90 \cdot 10^0$ | 1.78 | 33 | $1.36 \cdot 10^9$ | $5.20 \cdot 10^1$ | 4.74 | 335 |
| GRCh38 | 21 | 4 | 141,645,448 | $2.93 \cdot 10^7$ | $2.60 \cdot 10^7$ | $1.10 \cdot 10^0$ | 1.01 | 12 | $1.65 \cdot 10^8$ | $6.00 \cdot 10^0$ | 1.23 | 108 |
| GRCh38 | 27 | 4 | 107,171,949 | $2.93 \cdot 10^7$ | $2.82 \cdot 10^7$ | $1.10 \cdot 10^0$ | 1.01 | 12 | $1.91 \cdot 10^8$ | $6.00 \cdot 10^0$ | 1.25 | 66 |
| GRCh38 | 32 | 4 | 98,414,723 | $2.93 \cdot 10^7$ | $2.92 \cdot 10^7$ | $1.10 \cdot 10^0$ | 1.01 | 8 | $1.85 \cdot 10^8$ | $6.00 \cdot 10^0$ | 1.22 | 70 |
| SRR622461 | 15 | 4 | 117,393,511 | $7.90 \cdot 10^7$ | $4.93 \cdot 10^7$ | $3.30 \cdot 10^0$ | 2.2 | 55 | $1.34 \cdot 10^{10}$ | $9.60 \cdot 10^1$ | 20.68 | 398 |
| SRR622461 | 21 | 3 | 2,385,949,540 | $7.35 \cdot 10^7$ | $7.09 \cdot 10^7$ | $1.80 \cdot 10^0$ | 1.11 | 13 | $5.63 \cdot 10^8$ | $9.00 \cdot 10^0$ | 1.68 | 74 |
| SRR622461 | 27 | 4 | 1,782,384,901 | $6.80 \cdot 10^7$ | $4.92 \cdot 10^7$ | $1.30 \cdot 10^0$ | 1.04 | 6 | $4.58 \cdot 10^8$ | $8.00 \cdot 10^0$ | 1.53 | 21 |
| SRR622461 | 32 | 4 | 1,719,165,525 | $6.34 \cdot 10^7$ | $4.95 \cdot 10^7$ | $1.30 \cdot 10^0$ | 1.03 | 5 | $4.01 \cdot 10^8$ | $7.00 \cdot 10^0$ | 1.48 | 18 |

**Memory comparison for epsilon = 0.01**

**Table 2:**
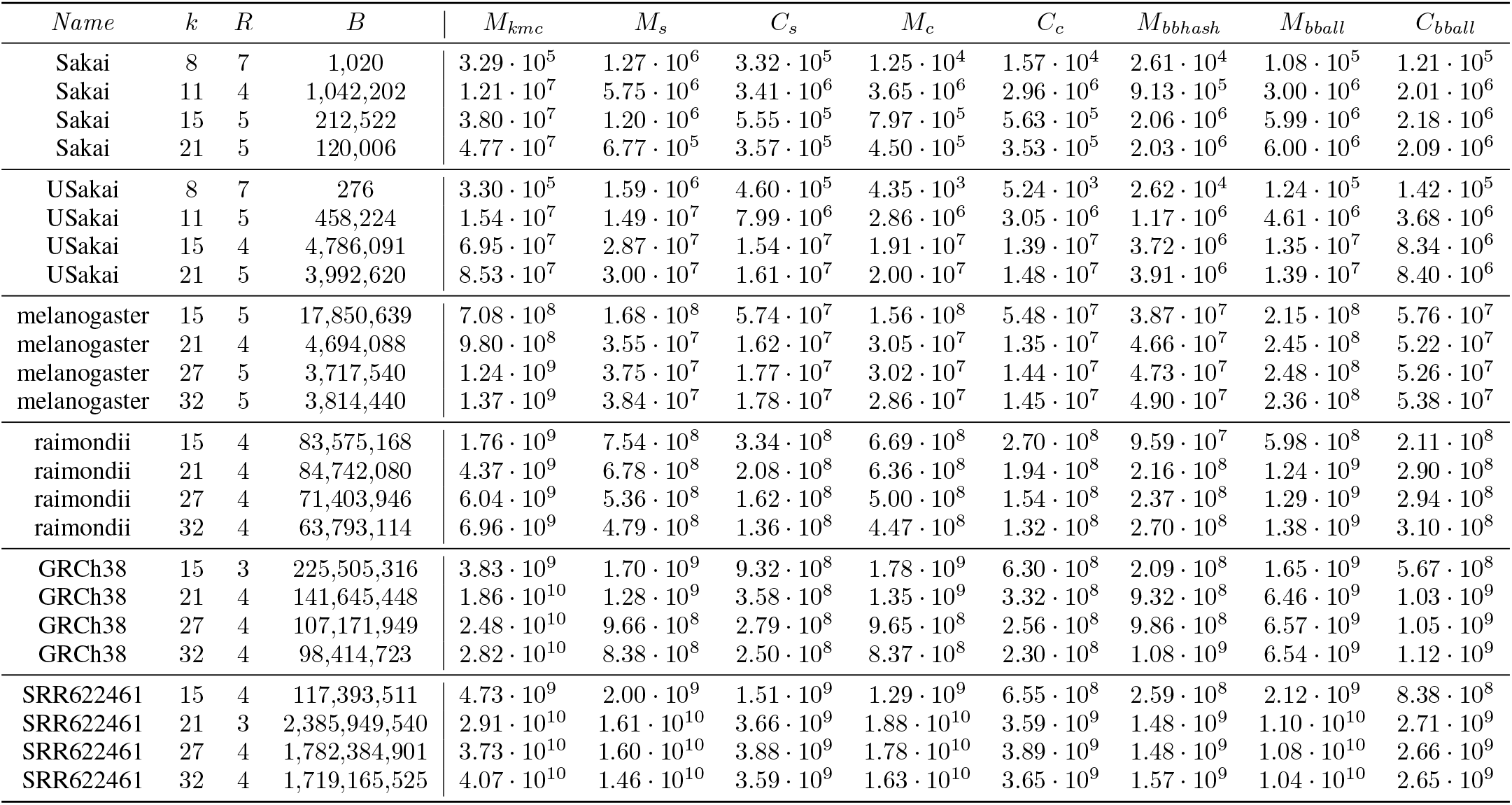
Full table comparing the memory of Set-Min sketch constructed for *E* = 0.01 to all other variables for all possible *k*s. *R* = number of rows, *B* = number of columns, *M*_*kmc*_, *M*_*s*_ = size of Set-Min sketch in bytes, *C*_*s*_ = compressed Set-Min sketch size, *M*_*c*_ = size of Count-Min sketch in bytes, *C*_*c*_ = compressed size of Count-Min sketch, *M*_*bbhash*_ = size of the MPHF generated by BBHash without the external array, *M*_*bball*_ = total memory for BBHash taking into account the external array, *C*_*bball*_ = compressed size of *M*_*bball*_

| *Name* | *k* | *R* | *B* | $M_{kmc}$ | $M_s$ | $C_s$ | $M_c$ | $C_c$ | $M_{bbhash}$ | $M_{bball}$ | $C_{bball}$ |
| --- | --- | --- | --- | --- | --- | --- | --- | --- | --- | --- | --- |
| Sakai | 8 | 7 | 1,020 | $3.29 \cdot 10^5$ | $1.27 \cdot 10^6$ | $3.32 \cdot 10^5$ | $1.25 \cdot 10^4$ | $1.57 \cdot 10^4$ | $2.61 \cdot 10^4$ | $1.08 \cdot 10^5$ | $1.21 \cdot 10^5$ |
| Sakai | 11 | 4 | 1,042,202 | $1.21 \cdot 10^7$ | $5.75 \cdot 10^6$ | $3.41 \cdot 10^6$ | $3.65 \cdot 10^6$ | $2.96 \cdot 10^6$ | $9.13 \cdot 10^5$ | $3.00 \cdot 10^6$ | $2.01 \cdot 10^6$ |
| Sakai | 15 | 5 | 212,522 | $3.80 \cdot 10^7$ | $1.20 \cdot 10^6$ | $5.55 \cdot 10^5$ | $7.97 \cdot 10^5$ | $5.63 \cdot 10^5$ | $2.06 \cdot 10^6$ | $5.99 \cdot 10^6$ | $2.18 \cdot 10^6$ |
| Sakai | 21 | 5 | 120,006 | $4.77 \cdot 10^7$ | $6.77 \cdot 10^5$ | $3.57 \cdot 10^5$ | $4.50 \cdot 10^5$ | $3.53 \cdot 10^5$ | $2.03 \cdot 10^6$ | $6.00 \cdot 10^6$ | $2.09 \cdot 10^6$ |
| USakai | 8 | 7 | 276 | $3.30 \cdot 10^5$ | $1.59 \cdot 10^6$ | $4.60 \cdot 10^5$ | $4.35 \cdot 10^3$ | $5.24 \cdot 10^3$ | $2.62 \cdot 10^4$ | $1.24 \cdot 10^5$ | $1.42 \cdot 10^5$ |
| USakai | 11 | 5 | 458,224 | $1.54 \cdot 10^7$ | $1.49 \cdot 10^7$ | $7.99 \cdot 10^6$ | $2.86 \cdot 10^6$ | $3.05 \cdot 10^6$ | $1.17 \cdot 10^6$ | $4.61 \cdot 10^6$ | $3.68 \cdot 10^6$ |
| USakai | 15 | 4 | 4,786,091 | $6.95 \cdot 10^7$ | $2.87 \cdot 10^7$ | $1.54 \cdot 10^7$ | $1.91 \cdot 10^7$ | $1.39 \cdot 10^7$ | $3.72 \cdot 10^6$ | $1.35 \cdot 10^7$ | $8.34 \cdot 10^6$ |
| USakai | 21 | 5 | 3,992,620 | $8.53 \cdot 10^7$ | $3.00 \cdot 10^7$ | $1.61 \cdot 10^7$ | $2.00 \cdot 10^7$ | $1.48 \cdot 10^7$ | $3.91 \cdot 10^6$ | $1.39 \cdot 10^7$ | $8.40 \cdot 10^6$ |
| melanogaster | 15 | 5 | 17,850,639 | $7.08 \cdot 10^8$ | $1.68 \cdot 10^8$ | $5.74 \cdot 10^7$ | $1.56 \cdot 10^8$ | $5.48 \cdot 10^7$ | $3.87 \cdot 10^7$ | $2.15 \cdot 10^8$ | $5.76 \cdot 10^7$ |
| melanogaster | 21 | 4 | 4,694,088 | $9.80 \cdot 10^8$ | $3.55 \cdot 10^7$ | $1.62 \cdot 10^7$ | $3.05 \cdot 10^7$ | $1.35 \cdot 10^7$ | $4.66 \cdot 10^7$ | $2.45 \cdot 10^8$ | $5.22 \cdot 10^7$ |
| melanogaster | 27 | 5 | 3,717,540 | $1.24 \cdot 10^9$ | $3.75 \cdot 10^7$ | $1.77 \cdot 10^7$ | $3.02 \cdot 10^7$ | $1.44 \cdot 10^7$ | $4.73 \cdot 10^7$ | $2.48 \cdot 10^8$ | $5.26 \cdot 10^7$ |
| melanogaster | 32 | 5 | 3,814,440 | $1.37 \cdot 10^9$ | $3.84 \cdot 10^7$ | $1.78 \cdot 10^7$ | $2.86 \cdot 10^7$ | $1.45 \cdot 10^7$ | $4.90 \cdot 10^7$ | $2.36 \cdot 10^8$ | $5.38 \cdot 10^7$ |
| raimondii | 15 | 4 | 83,575,168 | $1.76 \cdot 10^9$ | $7.54 \cdot 10^8$ | $3.34 \cdot 10^8$ | $6.69 \cdot 10^8$ | $2.70 \cdot 10^8$ | $9.59 \cdot 10^7$ | $5.98 \cdot 10^8$ | $2.11 \cdot 10^8$ |
| raimondii | 21 | 4 | 84,742,080 | $4.37 \cdot 10^9$ | $6.78 \cdot 10^8$ | $2.08 \cdot 10^8$ | $6.36 \cdot 10^8$ | $1.94 \cdot 10^8$ | $2.16 \cdot 10^8$ | $1.24 \cdot 10^9$ | $2.90 \cdot 10^8$ |
| raimondii | 27 | 4 | 71,403,946 | $6.04 \cdot 10^9$ | $5.36 \cdot 10^8$ | $1.62 \cdot 10^8$ | $5.00 \cdot 10^8$ | $1.54 \cdot 10^8$ | $2.37 \cdot 10^8$ | $1.29 \cdot 10^9$ | $2.94 \cdot 10^8$ |
| raimondii | 32 | 4 | 63,793,114 | $6.96 \cdot 10^9$ | $4.79 \cdot 10^8$ | $1.36 \cdot 10^8$ | $4.47 \cdot 10^8$ | $1.32 \cdot 10^8$ | $2.70 \cdot 10^8$ | $1.38 \cdot 10^9$ | $3.10 \cdot 10^8$ |
| GRCh38 | 15 | 3 | 225,505,316 | $3.83 \cdot 10^9$ | $1.70 \cdot 10^9$ | $9.32 \cdot 10^8$ | $1.78 \cdot 10^9$ | $6.30 \cdot 10^8$ | $2.09 \cdot 10^8$ | $1.65 \cdot 10^9$ | $5.67 \cdot 10^8$ |
| GRCh38 | 21 | 4 | 141,645,448 | $1.86 \cdot 10^{10}$ | $1.28 \cdot 10^9$ | $3.58 \cdot 10^8$ | $1.35 \cdot 10^9$ | $3.32 \cdot 10^8$ | $9.32 \cdot 10^8$ | $6.46 \cdot 10^9$ | $1.03 \cdot 10^9$ |
| GRCh38 | 27 | 4 | 107,171,949 | $2.48 \cdot 10^{10}$ | $9.66 \cdot 10^8$ | $2.79 \cdot 10^8$ | $9.65 \cdot 10^8$ | $2.56 \cdot 10^8$ | $9.86 \cdot 10^8$ | $6.57 \cdot 10^9$ | $1.05 \cdot 10^9$ |
| GRCh38 | 32 | 4 | 98,414,723 | $2.82 \cdot 10^{10}$ | $8.38 \cdot 10^8$ | $2.50 \cdot 10^8$ | $8.37 \cdot 10^8$ | $2.30 \cdot 10^8$ | $1.08 \cdot 10^9$ | $6.54 \cdot 10^9$ | $1.12 \cdot 10^9$ |
| SRR622461 | 15 | 4 | 117,393,511 | $4.73 \cdot 10^9$ | $2.00 \cdot 10^9$ | $1.51 \cdot 10^9$ | $1.29 \cdot 10^9$ | $6.55 \cdot 10^8$ | $2.59 \cdot 10^8$ | $2.12 \cdot 10^9$ | $8.38 \cdot 10^8$ |
| SRR622461 | 21 | 3 | 2,385,949,540 | $2.91 \cdot 10^{10}$ | $1.61 \cdot 10^{10}$ | $3.66 \cdot 10^9$ | $1.88 \cdot 10^{10}$ | $3.59 \cdot 10^9$ | $1.48 \cdot 10^9$ | $1.10 \cdot 10^{10}$ | $2.71 \cdot 10^9$ |
| SRR622461 | 27 | 4 | 1,782,384,901 | $3.73 \cdot 10^{10}$ | $1.60 \cdot 10^{10}$ | $3.88 \cdot 10^9$ | $1.78 \cdot 10^{10}$ | $3.89 \cdot 10^9$ | $1.48 \cdot 10^9$ | $1.08 \cdot 10^{10}$ | $2.66 \cdot 10^9$ |
| SRR622461 | 32 | 4 | 1,719,165,525 | $4.07 \cdot 10^{10}$ | $1.46 \cdot 10^{10}$ | $3.59 \cdot 10^9$ | $1.63 \cdot 10^{10}$ | $3.65 \cdot 10^9$ | $1.57 \cdot 10^9$ | $1.04 \cdot 10^{10}$ | $2.65 \cdot 10^9$ |

**Performance comparison for epsilon = 0.001**

**Table 3:** Full table comparing the performance of Set-Min sketch constructed for *E* = 0.001 to all other variables for all possible *k*s. *R* = number of rows, *B* = number of columns, *T* = theoretical maximum error bound, *E*_*s*_ = sum of errors for Set-Min, *N*_*s*_ = total number of collisions for Set-Min, *A*_*s*_ = average point-query error (*E*_*s*_*/N*_*s*_), 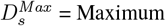 point-query error for Set-Min, *E*_*c*_ = sum of errors for Count-Min, *N*_*c*_ = total number of collisions for Count-Min, *A*_*c*_ = average point-query error (*E*_*c*_*/N*_*c*_), 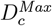 point-query error for Count-Min

| *Name* | *k* | *R* | *B* | *T* | *E<sub>s</sub>* | *N<sub>s</sub>* | *A<sub>s</sub>* | *D<sub>s</sub><sup>Max</sup>* | *E<sub>c</sub>* | *N<sub>c</sub>* | *A<sub>c</sub>* | *D<sub>c</sub><sup>Max</sup>* |
| --- | --- | --- | --- | --- | --- | --- | --- | --- | --- | --- | --- | --- |
| Sakai | 8 | 8 | 1,227 | $5.50 \cdot 10^3$ | 4,921 | $4.00 \cdot 10^{-1}$ | 21.03 | 76 | $2.24 \cdot 10^8$ | $1.00 \cdot 10^2$ | 3,429.11 | 4,933 |
| Sakai | 11 | 6 | 1,111,682 | $5.50 \cdot 10^3$ | 5,332 | $2.00 \cdot 10^{-1}$ | 1.05 | 3 | $4.78 \cdot 10^5$ | $1.10 \cdot 10^1$ | 1.85 | 13 |
| Sakai | 15 | 8 | 189,752 | $5.50 \cdot 10^3$ | 5,405 | $1.00 \cdot 10^{-1}$ | 1 | 1 | 60,364 | $1.00 \cdot 10^0$ | 1.05 | 4 |
| Sakai | 21 | 9 | 95,243 | $5.50 \cdot 10^3$ | 5,142 | $1.00 \cdot 10^{-1}$ | 1.01 | 2 | $1.69 \cdot 10^5$ | $3.00 \cdot 10^0$ | 1.07 | 5 |
| USakai | 8 | 10 | 236 | $2.62 \cdot 10^4$ | 28,302 | $5.00 \cdot 10^{-1}$ | 88.72 | 325 | $6.48 \cdot 10^9$ | $1.00 \cdot 10^2$ | 98,860.29 | $1.17 \cdot 10^5$ |
| USakai | 11 | 6 | 610,965 | $2.57 \cdot 10^4$ | 18,949 | $3.00 \cdot 10^{-1}$ | 2.12 | 14 | $4.18 \cdot 10^7$ | $9.40 \cdot 10^1$ | 14.58 | 81 |
| USakai | 15 | 6 | 5,105,164 | $2.49 \cdot 10^4$ | 23,817 | $2.00 \cdot 10^{-1}$ | 1.19 | 4 | $3.15 \cdot 10^6$ | $1.50 \cdot 10^1$ | 2.1 | 11 |
| USakai | 21 | 7 | 4,562,994 | $2.38 \cdot 10^4$ | 19,817 | $2.00 \cdot 10^{-1}$ | 1.12 | 3 | $2.88 \cdot 10^6$ | $1.50 \cdot 10^1$ | 1.96 | 11 |
| melanogaster | 15 | 6 | 24,792,555 | $1.43 \cdot 10^5$ | $1.32 \cdot 10^5$ | $1.00 \cdot 10^{-1}$ | 1 | 2 | $1.20 \cdot 10^6$ | $1.00 \cdot 10^0$ | 1.18 | 10 |
| melanogaster | 21 | 8 | 3,352,920 | $1.43 \cdot 10^5$ | $1.27 \cdot 10^5$ | $1.00 \cdot 10^{-1}$ | 1 | 4 | $9.23 \cdot 10^6$ | $6.00 \cdot 10^0$ | 1.23 | 19 |
| melanogaster | 27 | 8 | 3,319,232 | $1.43 \cdot 10^5$ | $1.29 \cdot 10^5$ | $1.00 \cdot 10^{-1}$ | 1 | 3 | $8.68 \cdot 10^6$ | $6.00 \cdot 10^0$ | 1.21 | 19 |
| melanogaster | 32 | 8 | 3,405,750 | $1.42 \cdot 10^5$ | $1.30 \cdot 10^5$ | $1.00 \cdot 10^{-1}$ | 1 | 3 | $7.83 \cdot 10^6$ | $5.00 \cdot 10^0$ | 1.19 | 17 |
| raimondii | 15 | 6 | 89,146,846 | $7.27 \cdot 10^5$ | $6.71 \cdot 10^5$ | $3.00 \cdot 10^{-1}$ | 1.02 | 5 | $4.99 \cdot 10^7$ | $1.10 \cdot 10^1$ | 1.77 | 24 |
| raimondii | 21 | 7 | 69,177,208 | $7.27 \cdot 10^5$ | $6.07 \cdot 10^5$ | $1.00 \cdot 10^{-1}$ | 1 | 3 | $1.12 \cdot 10^7$ | $2.00 \cdot 10^0$ | 1.15 | 11 |
| raimondii | 27 | 7 | 58,288,935 | $7.27 \cdot 10^5$ | $6.94 \cdot 10^5$ | $1.00 \cdot 10^{-1}$ | 1 | 3 | $8.89 \cdot 10^6$ | $1.00 \cdot 10^0$ | 1.11 | 9 |
| raimondii | 32 | 8 | 45,566,510 | $7.27 \cdot 10^5$ | $6.41 \cdot 10^5$ | $1.00 \cdot 10^{-1}$ | 1 | 2 | $7.73 \cdot 10^6$ | $1.00 \cdot 10^0$ | 1.08 | 7 |
| GRCh38 | 15 | 6 | 157,853,721 | $2.93 \cdot 10^6$ | $2.45 \cdot 10^6$ | $4.00 \cdot 10^{-1}$ | 1.24 | 9 | $1.18 \cdot 10^9$ | $5.50 \cdot 10^1$ | 3.95 | 58 |
| GRCh38 | 21 | 7 | 115,628,937 | $2.93 \cdot 10^6$ | $2.77 \cdot 10^6$ | $1.00 \cdot 10^{-1}$ | 1 | 3 | $4.42 \cdot 10^7$ | $2.00 \cdot 10^0$ | 1.09 | 14 |
| GRCh38 | 27 | 8 | 76,551,392 | $2.93 \cdot 10^6$ | $2.59 \cdot 10^6$ | $1.00 \cdot 10^{-1}$ | 1 | 2 | $5.81 \cdot 10^7$ | $2.00 \cdot 10^0$ | 1.08 | 13 |
| GRCh38 | 32 | 8 | 70,296,231 | $2.93 \cdot 10^6$ | $2.69 \cdot 10^6$ | $1.00 \cdot 10^{-1}$ | 1 | 2 | $5.46 \cdot 10^7$ | $2.00 \cdot 10^0$ | 1.07 | 10 |
| SRR622461 | 15 | 5 | 140,872,213 | $7.90 \cdot 10^6$ | $7.38 \cdot 10^6$ | $6.00 \cdot 10^{-1}$ | 1.69 | 28 | $8.27 \cdot 10^9$ | $9.00 \cdot 10^1$ | 13.66 | 231 |

**Memory comparison for epsilon = 0.01**

**Table 4:**
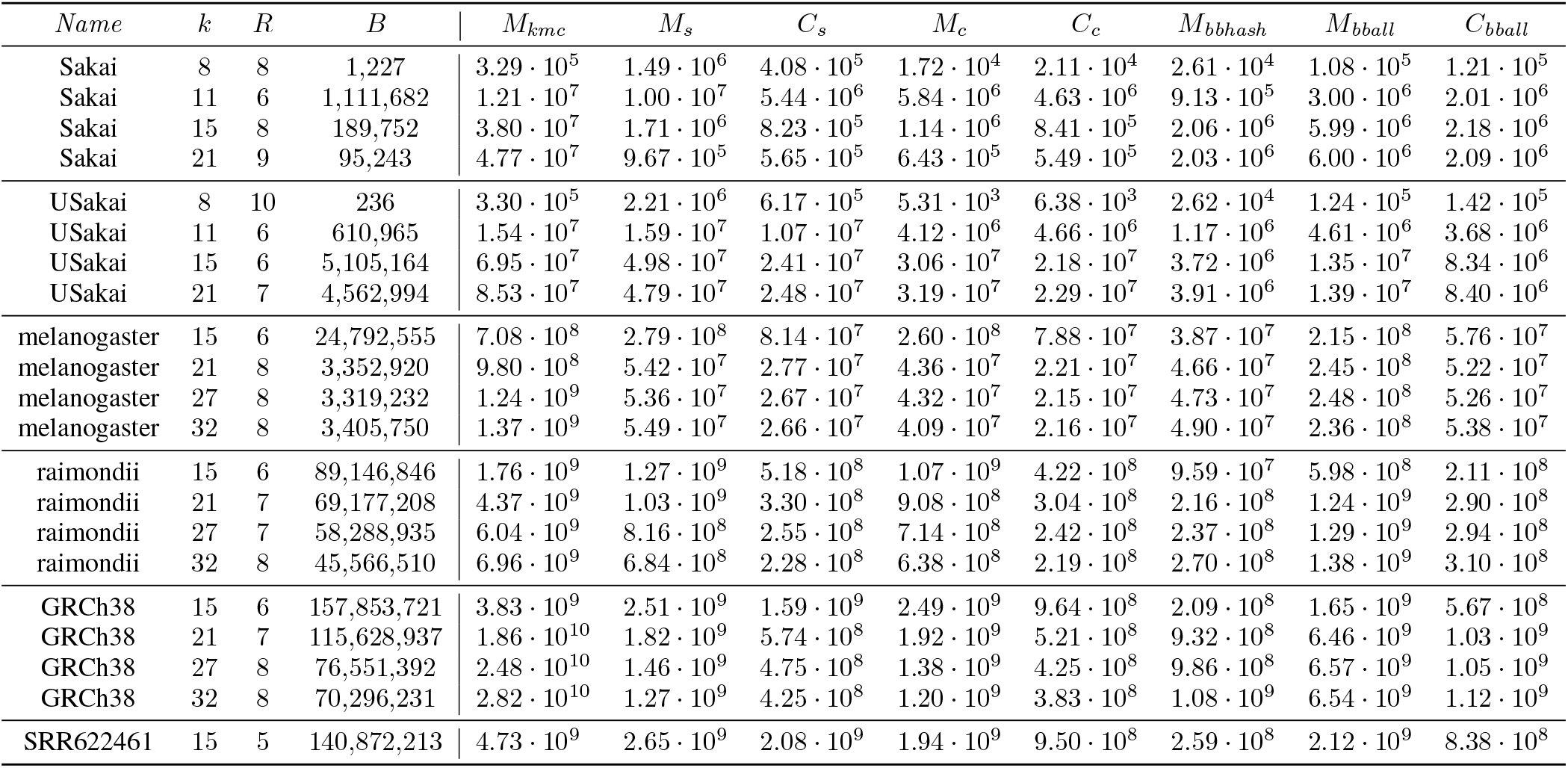
Full table comparing the memory of Set-Min sketch constructed for *E* = 0.001 to all other variables for all possible *k*s. *R* = number of rows, *B* = number of columns, *M*_*kmc*_, *M*_*s*_ = size of Set-Min sketch in bytes, *C*_*s*_ = compressed Set-Min sketch size, *M*_*c*_ = size of Count-Min sketch in bytes, *C*_*c*_ = compressed size of Count-Min sketch, *M*_*bbhash*_ = size of the MPHF generated by BBHash without the external array, *M*_*bball*_ = total memory for BBHash taking into account the external array, *C*_*bball*_ = compressed size of *M*_*bball*_

| *Name* | *k* | *R* | *B* | $M_{kmc}$ | $M_s$ | $C_s$ | $M_c$ | $C_c$ | $M_{bbhash}$ | $M_{bball}$ | $C_{bball}$ |
| --- | --- | --- | --- | --- | --- | --- | --- | --- | --- | --- | --- |
| Sakai | 8 | 8 | 1,227 | $3.29 \cdot 10^5$ | $1.49 \cdot 10^6$ | $4.08 \cdot 10^5$ | $1.72 \cdot 10^4$ | $2.11 \cdot 10^4$ | $2.61 \cdot 10^4$ | $1.08 \cdot 10^5$ | $1.21 \cdot 10^5$ |
| Sakai | 11 | 6 | 1,111,682 | $1.21 \cdot 10^7$ | $1.00 \cdot 10^7$ | $5.44 \cdot 10^6$ | $5.84 \cdot 10^6$ | $4.63 \cdot 10^6$ | $9.13 \cdot 10^5$ | $3.00 \cdot 10^6$ | $2.01 \cdot 10^6$ |
| Sakai | 15 | 8 | 189,752 | $3.80 \cdot 10^7$ | $1.71 \cdot 10^6$ | $8.23 \cdot 10^5$ | $1.14 \cdot 10^6$ | $8.41 \cdot 10^5$ | $2.06 \cdot 10^6$ | $5.99 \cdot 10^6$ | $2.18 \cdot 10^6$ |
| Sakai | 21 | 9 | 95,243 | $4.77 \cdot 10^7$ | $9.67 \cdot 10^5$ | $5.65 \cdot 10^5$ | $6.43 \cdot 10^5$ | $5.49 \cdot 10^5$ | $2.03 \cdot 10^6$ | $6.00 \cdot 10^6$ | $2.09 \cdot 10^6$ |
| USakai | 8 | 10 | 236 | $3.30 \cdot 10^5$ | $2.21 \cdot 10^6$ | $6.17 \cdot 10^5$ | $5.31 \cdot 10^3$ | $6.38 \cdot 10^3$ | $2.62 \cdot 10^4$ | $1.24 \cdot 10^5$ | $1.42 \cdot 10^5$ |
| USakai | 11 | 6 | 610,965 | $1.54 \cdot 10^7$ | $1.59 \cdot 10^7$ | $1.07 \cdot 10^7$ | $4.12 \cdot 10^6$ | $4.66 \cdot 10^6$ | $1.17 \cdot 10^6$ | $4.61 \cdot 10^6$ | $3.68 \cdot 10^6$ |
| USakai | 15 | 6 | 5,105,164 | $6.95 \cdot 10^7$ | $4.98 \cdot 10^7$ | $2.41 \cdot 10^7$ | $3.06 \cdot 10^7$ | $2.18 \cdot 10^7$ | $3.72 \cdot 10^6$ | $1.35 \cdot 10^7$ | $8.34 \cdot 10^6$ |
| USakai | 21 | 7 | 4,562,994 | $8.53 \cdot 10^7$ | $4.79 \cdot 10^7$ | $2.48 \cdot 10^7$ | $3.19 \cdot 10^7$ | $2.29 \cdot 10^7$ | $3.91 \cdot 10^6$ | $1.39 \cdot 10^7$ | $8.40 \cdot 10^6$ |
| melanogaster | 15 | 6 | 24,792,555 | $7.08 \cdot 10^8$ | $2.79 \cdot 10^8$ | $8.14 \cdot 10^7$ | $2.60 \cdot 10^8$ | $7.88 \cdot 10^7$ | $3.87 \cdot 10^7$ | $2.15 \cdot 10^8$ | $5.76 \cdot 10^7$ |
| melanogaster | 21 | 8 | 3,352,920 | $9.80 \cdot 10^8$ | $5.42 \cdot 10^7$ | $2.77 \cdot 10^7$ | $4.36 \cdot 10^7$ | $2.21 \cdot 10^7$ | $4.66 \cdot 10^7$ | $2.45 \cdot 10^8$ | $5.22 \cdot 10^7$ |
| melanogaster | 27 | 8 | 3,319,232 | $1.24 \cdot 10^9$ | $5.36 \cdot 10^7$ | $2.67 \cdot 10^7$ | $4.32 \cdot 10^7$ | $2.15 \cdot 10^7$ | $4.73 \cdot 10^7$ | $2.48 \cdot 10^8$ | $5.26 \cdot 10^7$ |
| melanogaster | 32 | 8 | 3,405,750 | $1.37 \cdot 10^9$ | $5.49 \cdot 10^7$ | $2.66 \cdot 10^7$ | $4.09 \cdot 10^7$ | $2.16 \cdot 10^7$ | $4.90 \cdot 10^7$ | $2.36 \cdot 10^8$ | $5.38 \cdot 10^7$ |
| raimondii | 15 | 6 | 89,146,846 | $1.76 \cdot 10^9$ | $1.27 \cdot 10^9$ | $5.18 \cdot 10^8$ | $1.07 \cdot 10^9$ | $4.22 \cdot 10^8$ | $9.59 \cdot 10^7$ | $5.98 \cdot 10^8$ | $2.11 \cdot 10^8$ |
| raimondii | 21 | 7 | 69,177,208 | $4.37 \cdot 10^9$ | $1.03 \cdot 10^9$ | $3.30 \cdot 10^8$ | $9.08 \cdot 10^8$ | $3.04 \cdot 10^8$ | $2.16 \cdot 10^8$ | $1.24 \cdot 10^9$ | $2.90 \cdot 10^8$ |
| raimondii | 27 | 7 | 58,288,935 | $6.04 \cdot 10^9$ | $8.16 \cdot 10^8$ | $2.55 \cdot 10^8$ | $7.14 \cdot 10^8$ | $2.42 \cdot 10^8$ | $2.37 \cdot 10^8$ | $1.29 \cdot 10^9$ | $2.94 \cdot 10^8$ |
| raimondii | 32 | 8 | 45,566,510 | $6.96 \cdot 10^9$ | $6.84 \cdot 10^8$ | $2.28 \cdot 10^8$ | $6.38 \cdot 10^8$ | $2.19 \cdot 10^8$ | $2.70 \cdot 10^8$ | $1.38 \cdot 10^9$ | $3.10 \cdot 10^8$ |
| GRCh38 | 15 | 6 | 157,853,721 | $3.83 \cdot 10^9$ | $2.51 \cdot 10^9$ | $1.59 \cdot 10^9$ | $2.49 \cdot 10^9$ | $9.64 \cdot 10^8$ | $2.09 \cdot 10^8$ | $1.65 \cdot 10^9$ | $5.67 \cdot 10^8$ |
| GRCh38 | 21 | 7 | 115,628,937 | $1.86 \cdot 10^{10}$ | $1.82 \cdot 10^9$ | $5.74 \cdot 10^8$ | $1.92 \cdot 10^9$ | $5.21 \cdot 10^8$ | $9.32 \cdot 10^8$ | $6.46 \cdot 10^9$ | $1.03 \cdot 10^9$ |
| GRCh38 | 27 | 8 | 76,551,392 | $2.48 \cdot 10^{10}$ | $1.46 \cdot 10^9$ | $4.75 \cdot 10^8$ | $1.38 \cdot 10^9$ | $4.25 \cdot 10^8$ | $9.86 \cdot 10^8$ | $6.57 \cdot 10^9$ | $1.05 \cdot 10^9$ |
| GRCh38 | 32 | 8 | 70,296,231 | $2.82 \cdot 10^{10}$ | $1.27 \cdot 10^9$ | $4.25 \cdot 10^8$ | $1.20 \cdot 10^9$ | $3.83 \cdot 10^8$ | $1.08 \cdot 10^9$ | $6.54 \cdot 10^9$ | $1.12 \cdot 10^9$ |
| SRR622461 | 15 | 5 | 140,872,213 | $4.73 \cdot 10^9$ | $2.65 \cdot 10^9$ | $2.08 \cdot 10^9$ | $1.94 \cdot 10^9$ | $9.50 \cdot 10^8$ | $2.59 \cdot 10^8$ | $2.12 \cdot 10^9$ | $8.38 \cdot 10^8$ |

## Footnotes

1 Here we assume *k* to be sufficiently large, typically *k >* log_4_ *L*, where *L* is the data size.

2 http://flybase.org

3 ftp://ftp.ncbi.nlm.nih.gov/genomes/all/GCA/000/001/405/GCA_000001405.15_GRCh38/seqs_for_alignment_pipelines.ucsc_ids/GCA_000001405.

4 ftp://ftp.sra.ebi.ac.uk/vol1/fastq/SRR622/SRR622461/SRR622461_1.fastq.gz. Only the file SRR622461_1 is used in this study and we then omit the underscored nomenclature to simplify the notation.

